# The human circular Tau 12->7 RNA is translated in mouse neurons and promotes tau aggregation

**DOI:** 10.64898/2026.10.26.761947

**Authors:** Naghme Bagheri, Weikang Cai, Salim El-Amouri, Marc Dobler, Cole J. Cochran, Andrew K. Turner, Anirudhya Lahiri, Shuling Fister, David W. Rodgers, Yuan Wen, Peter T. Nelson, Stefan Stamm

## Abstract

In addition to numerous mRNA isoforms, the human microtubule associated protein tau (*MAPT*) gene generates circular RNAs that are not known to be expressed in rodents. Using adeno associated viruses, we expressed human circular Tau RNA generated from backsplicing of exon 12 to exon 7 (circTau 12->7) in mouse brain. We observed rolling circle translation of circTau 12->7, similar to cultured human HEK293 cells. Despite using a ubiquitously active promoter, circTau 12->7 protein and a control circEGFP protein were only observed in mouse neurons, indicating neuron-specific translation of these circular RNAs. CircTau 12->7 expression resulted in tau aggregates resembling neurofibrillary tangles in mouse brain that included endogenous mouse tau protein. Our data support a model where neuron-specific translation of circTau RNAs induces ‘linear’ mRNA-translated tau aggregation, which could contribute to tauopathies, including Alzheimer’s disease. Thus, targeting circTau RNAs could be a therapeutic approach for Alzheimer’s disease.

## Introduction

Circular RNAs (circRNAs) are covalently closed RNAs, mostly generated through backsplicing of a 5′ splice site to a downstream 3′ splice site (1). CircRNAs are present in cytoplasm, are highly expressed in brain (2–5), and are enriched in synaptosomes (6). In a recent study, the expression levels of 14 circRNAs were associated with Braak neurofibrillary tangle (NFT) stages in Alzheimer’s disease (AD), and explained 31% of the clinical dementia rating (7).

There is increasing evidence that circRNAs can be translated (8), especially under conditions where 5′ methylguanosine cap-dependent translation is inhibited. We discovered human-specific circRNAs transcribed from the microtubule associated protein tau (9) (*MAPT*) gene, and showed that these circularized tau RNAs (circTau) can be translated after adenosine to inosine circRNA modification (10).

The tau protein stabilizes microtubules (11) through binding to tubulin via three or four microtubule binding repeat motifs (numbered R1-R4), formed by *MAPT* exons 7-12. In dozens of human diseases known as tauopathies, tau protein misfolds into intracellular, insoluble protein polymers (12). These pathologic polymers aggregate into intraneuronal tau inclusions (neurofibrillary tangles, NFTs) in AD, that are the pathological feature most strongly associated with cognitive impairment (13,14).

We previously showed that circTau-encoded proteins can promote the aggregation of ‘linear’ mRNA encoded tau protein *in vitro* and in reporter cells (15), which might reflect the general property of circRNA-encoded proteins to promote pathogenetic phase separation and protein aggregation (16).

In the current study, to test the effect of circTau RNAs under physiological conditions in mammalian brain, we generated adeno associated viruses (AAVs) expressing the previously studied circTau 12->7 RNA. This circTau RNA is generated by backsplicing exon 12 to exon 7, creating a circular RNA of 681 nt in length (9). CircTau 12->7 contains only one start codon and no stop codons, and generates an indefinite open reading frame that encodes repetitions of tau peptides. These peptides, in turn, correspond to the fragments of the mRNA-encoded, ‘linear’ tau protein that contain the microtubule binding sites R1-R4. The repeated tau peptide fragments correspond to the K18 peptide that can cause tau aggregation (17). We here analyzed circTau 12->7 in several contexts. We found that circTau 12->7 RNA is present in polysomes in HEK293 cells, and upon injection into mouse brain, the circRNA is translated into proteins that reflect a rolling circle translation. AAV experiments indicated that the circTau 12->7 encoded protein interacts with constitutively expressed (mRNA-encoded) mouse tau protein and cause its aggregation into structures that histopathologically resemble NFTs.

## Material and Methods

### Transfection into cultured human (HEK293T) cells

cDNA transfections was performed in human embryonic kidney (HEK293T) (ATCC, CRL-3216) cells using the PEI Prime™ 1mg/ml (SeroChem, SKU: Prime-AQ100-5ml) transfection protocol. Cells were seeded in the DMEM (Gibco™ DMEM high glucose, Fisher Scientific, Catalog No. 11965-092) containing 10% fetal bovine serum (GeminiBio, CAT: 100-800) in 100-mm culture dishes (VWR, Cat.No.10062-880) to 50% confluency on the day of transfection. On the day of transfection, the medium was replaced with reduced-serum medium containing 1% FBS. 1-2 μg of DNA and 2-4 μg of PEI were used per 10⁶ cells. DNA and PEI were mixed in culture media with 1% FBS incubated at room temperature for 10 minutes and then added gradually to the cells. Cells were collected 24 hours after transfection for RNA extraction and 3 days after transfection for protein extraction.

### Immunoprecipitation

Transfected cells were washed with 1xPBS (Invitrogen by Thermo Fisher Scientific, AM9625), and lysed using low salt RIPA buffer (50 mM Tris-HCl (pH 7.4), 150 mM NaCl, 1% NP-40 with Protease Inhibitor Cocktail tablet (Milipore Sigma, 11836170001)), The lysates were boiled at 95° C for 4 minutes. Lysates were precleared using 20 μl Mouse IgG Coupled 4% agarose beads (9.4 IgG mg/ml) (Aviva System Biology, OOIB00007) and Benzonase (Millipore Sigma,≥250units/ μl ) in the presence of 1mM MgCl2. Precleared lysates were immunoprecipitated by 30 μl Anti FLAG M2 magnetic Bead (Miliore Sigma,M8823). Lysate and immunoprecipitate samples were used for western blot. Anti flag mouse Antibody (Millipore Sigma, A8592) was used to detect proteins.

### Polysome preparation

HEK293T cells were seeded in 150-mm dishes (two dishes per condition) and transfected at 30–50% confluency using polyethylenimine (PEI) in DMEM supplemented with 1% FBS. Transfections for polysomes were performed using 14 µg total DNA at a reporter:ADAR or a plasmid expressing enhanced green fluorescent protein (pEGFP) with plasmid ratio of 10.5:3.5 µg and 24 µg PEI. DNA complexes were prepared in DMEM, incubated for 10 min at room temperature, and added to the cells. Twenty-four hours after transfection, the medium was replaced with complete growth medium containing 10% FBS, and cells were incubated for an additional 2 h. To arrest translation and stabilize polysomes, cells were treated with cycloheximide (CHX; 100 µg/mL final concentration) for 10 min at 37°C. Cells were washed twice with ice-cold PBS containing CHX, harvested, and pooled from two plates per condition. Following centrifugation (300 × g, 10 min, 4°C), cell pellets were resuspended in 500 µL ice-cold polysome lysis buffer containing 20 mM Tris-HCl (pH 7.5), 100 mM KCl, 5 mM MgCl₂, 1% Triton X-100, 25X cOmplete™ Mini EDTA-free Protease Inhibitor Cocktail, 1 U/µL recombinant RNasin ribonuclease inhibitor, and 100 µg/mL CHX. Lysates were homogenized by pipetting, incubated on ice for 15 min with intermittent vortexing, and processed for downstream polysome isolation.

### Polysome isolation

Equal amounts of polysome lysate (≤1 mL) were layered onto 10–50% sucrose gradients and centrifuged in an SW41Ti swinging-bucket rotor at 39,000 rpm for 90 min at 4°C using maximum acceleration and brake settings. Following centrifugation, gradients were maintained on ice and fractionated using a Biocomp piston gradient fractionator. Fractions were collected at a flow rate of 1 mL/min, yielding 18 fractions per gradient.

### Adeno associated viruses (AAV)

The AAVs were commercially packaged by Vectorbuilder, Inc, into the C9 serotype. As controls, we used AAVs expressing circular EGFP and AAVs expressing standard, mRNA-encoded EGFP (pAAV[Exp]-CAG>EGFP:WPRE Vector builder AAV9M (VB 010000-9287ffw). To control for circRNA-specific effects, we created an expression construct where EGFP is split at the chromophore and the expression cassette flanked with HIPK3 introns (Supplemental Data 4). The constructs pAAV12-7-HIPK3 and pAAV-circEGFP-HIPK3 have been deposited into Addgene.

### Stereotactic injection and tissue processing

3-month-old male C57BL/6J mice were anesthetized with intraperitoneal injection of a ketamine (10 mg/ml) and xylazine (1 mg/ml) mix and placed onto a stereotaxic station (Stoelting). The skull was exposed via a small incision, and a small hole was drilled to desired coordinates. AAV9 virus (0.5 µl, 0.1 µl/min) encoding circTau 12->7, circEGFP, or regular EGFP were injected into the hippocampus (AP -2.0 mm; ML -1.5 mm; DV -2.0 mm from dura). Two to four weeks after injection, mice were anesthetized, transcardially perfused with 1x PBS (18). For immunohistochemical studies, mouse brains were extracted and fixed in 10% buffered formalin, followed by alcohol dehydration, paraffin embedding, and tissue sectioning. For biochemical and molecular studies, hippocampi from injected side and control side of each mouse were microdissected and snap-frozen in liquid nitrogen for RNA and protein analysis.

### RT-PCR

Cells were collected 24 hours after transfection, and total RNA was isolated using the PureLink™ RNA Mini Kit (Invitrogen; Cat. No. 12183025) according to the manufacturer’s instructions. Circular RNA was enriched using RNAseR (Abcam, 5 U, per reaction), followed by phenol-chloroform extraction. One-step RT-PCR was performed using the SuperScript™ IV UniPrime One-Step RT-PCR System (Invitrogen; REF 12596025). Reverse transcription was carried out at 50 °C for 10 minutes, followed by RT inactivation/ initial denaturation at 98 °C for 2 minutes. PCR amplification was performed for 30 cycles, followed by a final extension at 72 °C for 5 minutes. The primers used for amplification circTau12-7 for 3: CAGTCGAAGATTGGGTCCCT circTau12-7 rev 3: TGGAATCCTGGTGGCGTTG

### Regents for immunohistochemistry

Anti GFP (SYSY, guinea pig, polyclonal), anti-NeuN (D4G4O) Rabbit Monoclonal, (Cell Signalling), anti-FLAG (NB600-344) (Novus), anti-tau chicken, AB_2313563, polyclonal, (AvesLab), anti-tau, AF3494 goat, (R&DSystems), phosphor Tau AT8, Thermofisher, detection was with ABC Vectastain HRP kits (Vector Lab).

## Results

### circTau 12-> 7 RNA can be expressed in cultured human cells when flanked by short HIPK3 introns

We previously showed that circTau12->7 undergoes backsplicing when the cDNA reflecting the circular RNA is flanked by either the authentic tau introns or heterologous ZKSCAN introns (10). As our circTau12->7 expressing constructs (10,19) were too large to be used in AAVs, we exchanged the flanking introns with shortened HIPK3 introns known to promote RNA circularization (20). We inserted in frame stop codons in the 5’ intron upstream of the circTau12->7 backsplicing cassette, preventing translation of linear pre-mRNAs in any frame. The only start codon for an open reading frame present in the construct is downstream of the FLAG tag, ensuring that only protein made from the circular RNA contains the 3x FLAG tag (**Figure 1A**). Together with the FLAG tag, the circTau 12->7 RNA encompasses 762 nucleotides, a number divisible by three. It contains a single start codon and lacks an in-frame stop codon (**Supplemental Figure 1**). The circTau RNA can thus undergo rolling circle translation that generates multimers of mRNA-encoded tau fragments, each of which contain the four microtubule binding repeats (**Figure 1B**). These constructs were transfected into HEK293 cells with or without DNA constructs expressing ADAR1-p150 (adenosine deaminase acting on RNA). We previously showed that ADAR activity strongly promotes translation of the circTau 12->7 and circTau 12->10 RNAs (10). Here we first tested circRNA expression using total RNA and circRNA-specific primers that flank the backsplice site. The PCR resulted in the expected band of 228 nt, which was absent in controls (**Figure1 C, Supplemental Figure 1**). Both the direction of the PCR primers spanning the backsplice junction and the RNAse R resistance of the signal indicate that the amplicon was generated from a circular RNA. We next tested circTau 12->7 encoded protein expression after three days of transfection with or without co-transfected ADAR1-p150. CircTau 12->7 protein is heat stable and as in previous studies (10), we employed FLAG-immunoprecipitation from boiled lysates. We detected circTau 12->7 protein expression that was enhanced in the presence of ADAR1-p150, as observed in earlier studies (10). We observed protein bands on immunoblots with apparent length of 33, 65, and 112 kDa, which reflects the rolling circle translation. Each round of translation from start to start codon encoded a protein of 27 kDa, and the stop sites observed during rolling circle translation likely indicate ribosome stalling at defined circRNA regions. These data indicated that circTau 12->7 RNA was generated when flanked by heterologous HIPK3 introns. Similar to circTau 12->7 RNA flanked by natural and ZKSCAN introns (10), this RNA was translated, and that translation was strongly increased by ADAR activity.

**Figure 1:**
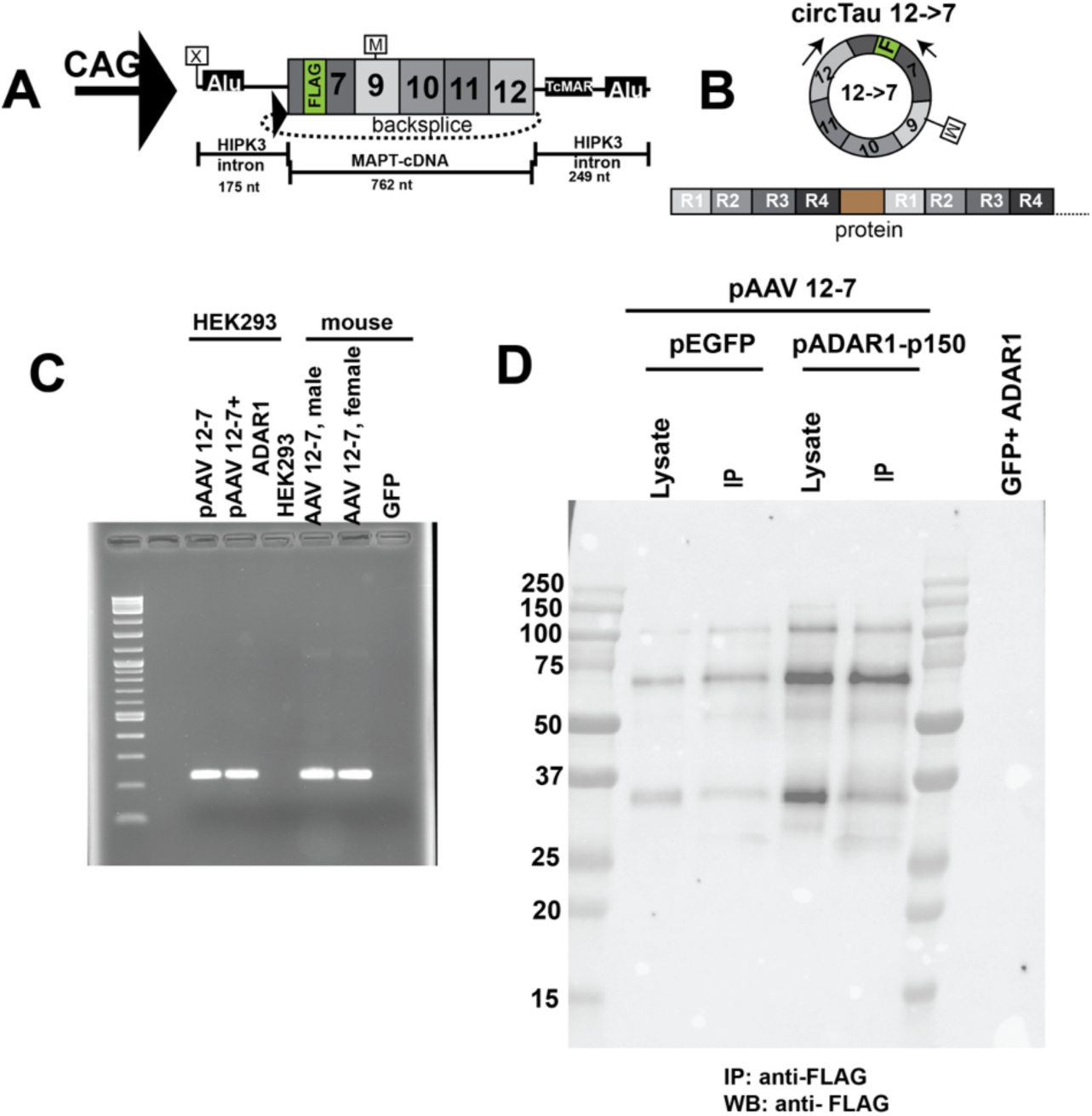
Heterologous HIPK3 introns flanking Tau 7->12 cDNA promote circularization in cultured HEK293 cells. **A. Structure of the construct pAAV circTau 12-7**. The circTau 12->7 expression cassette was cloned between HIPK3 heterologous intron, driven by a CAG promoter. The sequence of the insert is shown in Supplemental Figure 1. **B. circTau12->7 circular RNA and predicted proteins.** Backsplicing from exon 12 to 7 generates the circTau 12->7 RNA that contains one in-frame start codon in exon 7 and no stop codon, resulting in rolling circle translation. The resulting protein is a multimer of the mRNA-encoded fragment containing the four microtubule binding sites. **C. Expression of circTau 12-7 RNA.** The circRNA expressing plasmid pAAV-circTau12-7 was transfected into HEK293 cells in the presence and absence of ADAR1-p150 expression constructs. In parallel AAV-C9 viruses generated from this construct were injected into mouse hippocampus and total RNA was isolated from infected regions. Total RNA was treated with RNAse R before amplified using backsplice junction spanning primers (Supplemental Figure 1). **D. CircTau 12-7 RNA is translated in HEK293 cells**. The circTau 12-7 RNA expression construct pAAV 12-7 was transfected into HEK293 cells using lipofectamine. After 3 days, cells were lysed, lysates boiled and subjected to anti FLAG-immunoprecipitation. Protein was detected with anti-FLAG antibody.

### CircTau 12->7 RNA is present in polysomes

The association of circTau RNA with ribosomes is a necessary condition for circTau 12->7 RNA to be translated. We thus performed polysome analysis from HEK293 cells transfected with circTau 12-7 expression constructs. After 24 hrs of transfection, translation was arrested using cycloheximide, and polysomes were separated on 10 to 50% sucrose gradients (**Figure 2**). We detected circTau 12->7 RNA in RNAseR treated samples and observed a strong association with the 80S monosomes and a decreasing signal in larger polysomes (**Figure 2A**). When we cotransfected cDNAs expressing ADAR1-p150 with circTau12->7 expression clones, we observed a marked shift of circTau RNA towards larger polysomes (**Fig 2B**). There was no signal in GFP transfected cells, demonstrating the absence of intrinsic/constitutively expressed circTau12->7 RNAs in control-transfected HEK293 cells (**Figure 2C**). The circTau RNA signal was resistant to RNAse R, demonstrating the circular nature of the RNA, a result that apparently rules out a mRNA generated by trans-splicing. Instead, the data suggest that the RNA and protein signal we see is generated from translated circRNAs. Reflecting the increase in translation, ADAR activity shifts circTau 12->7 RNA into larger polysomes, possibly by de-repressing stalled circular RNA in the 80S complex.

**Figure 2:**
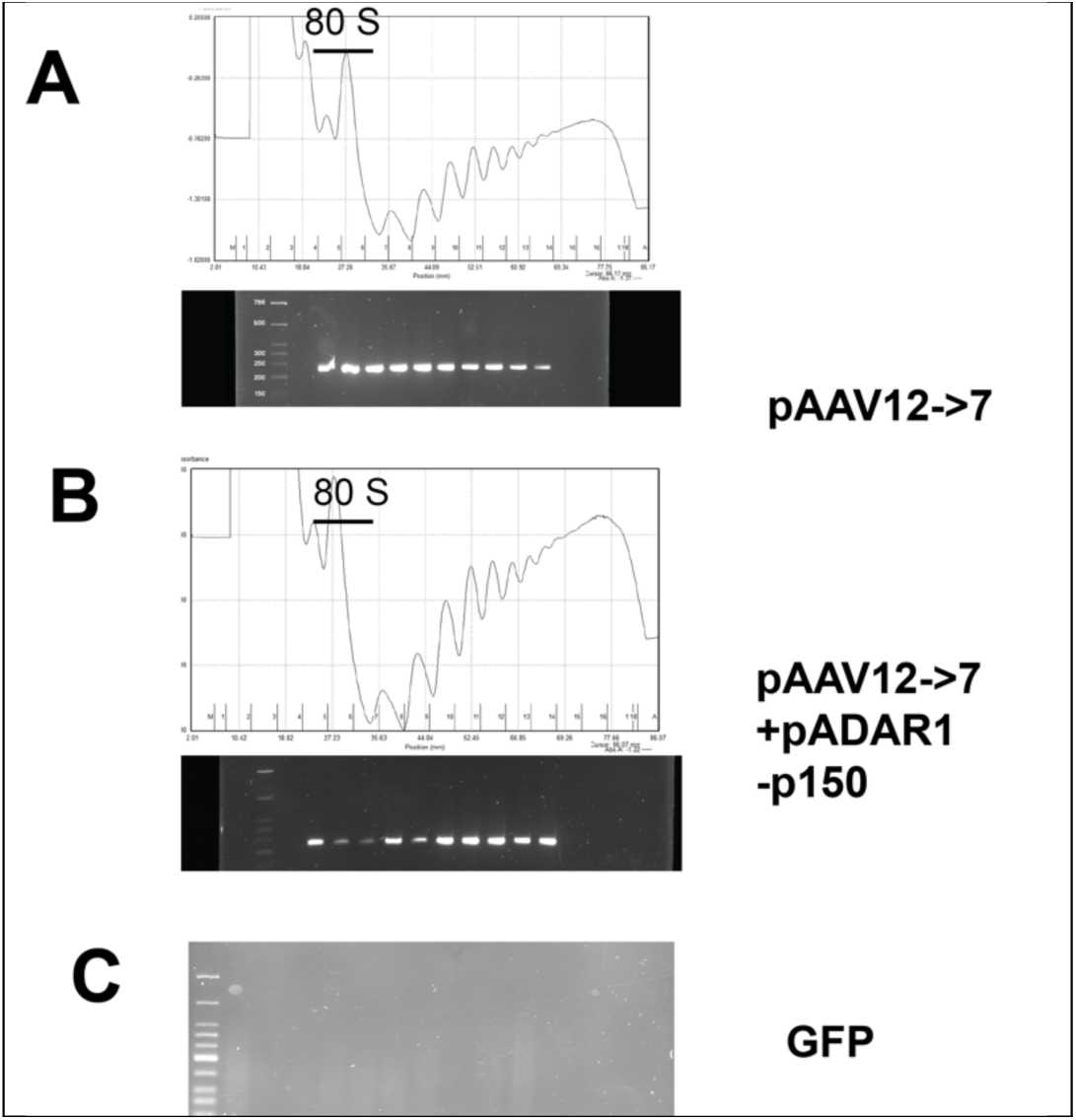
CircRNAs are present in polysomes. **A.** Polysome profile of HEK293 cells transfected with circTau 12->7 separated on 10-50% sucrose gradients. Total RNA was isolated from fractions and circRNAs were enriched through RNAseR treatment followed by RT-PCR using primers detecting circTau12->7. **B.** The circTau 12-> 7 expression clone was cotransfected with a clone expressing ADAR1-p150 and polysomes were generated as in (A). **C.** Amplification of GFP as negative control in the polysome fraction of A.

### CircTau 12->7 delivered by AAVs is expressed and translated in mouse brain

We next tested the expression of circTau 12->7 RNA in mouse brain after viral transduction. The circTau 12->7 expression construct (**Figure 1A**) was packaged into AAVs (serotype 9) with a promoter that has been shown to drive protein expression in multiple cell types (*i.e*., neurons and non-neuronal cells), including astrocytes (21). Five million GC (genomic copies) of the virus were delivered through intracranial injection into the hippocampus. We did not observe adverse behavioral effects after injection (data not shown). Animals were sacrificed after 21 days, at which timepoint hippocampi were microdissected and RNA analyzed by RT-PCR. RNA from areas injected with AAVs resulted in the expected amplicon, indicating that circTau 12->7 RNA is formed in mouse brain, similar to transfection in cultured human HEK293 cells (**Figure. 1C**).

The circTau12->7-encoded protein was analyzed by Western blot using immunoprecipates and total lysates from micro dissected injected areas. To allow comparisons, we kept the ratio between wet tissue and lysis buffer constant. We observed expression of proteins with apparent sizes of 33, 65, 112 and 150 kDa expression – a banding pattern quite similar to that after transfection in cultured human HEK293 cells. There was no difference in expression levels or apparent protein molecular weights between male and female mice (**Figure 3A**). Despite the overarching similarities, the protein expression in mice was much stronger than in HEK293 cells and the largest band of 150 kDa was visible in mice, but very faint in HEK293 cells. The pattern indicates a rolling circle translation in both systems, adding a peptide of 32, 47 and 38 kDa to each round of translation, which roughly reflects the 29 kDa for each round (**Figure 3A**). In summary, circTau RNAs are translated in mouse brain when delivered via AAVs.

**Figure 3:**
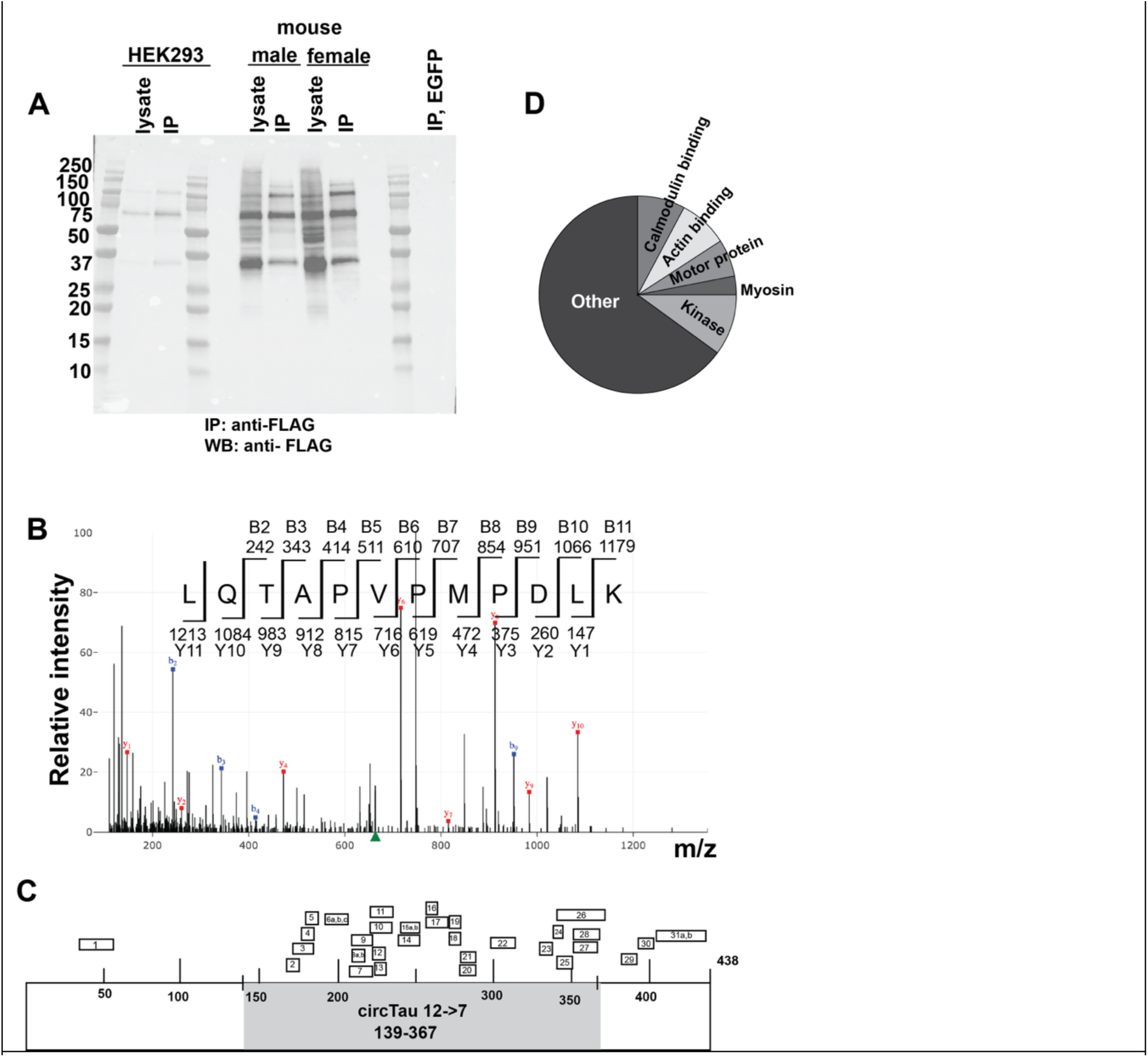
Translation of circTau 12->7 in mouse brain. **A. CircTau 12->7 protein expression in mouse brain.** BL6 mice were injected into the hippocampus with an AAV-C9 expressing the circTau 12->7 RNA or EGFP. After 21 days, injected areas were microdissected and analyzed by immunoprecipitations followed by Western blot. Lysates and immunoprecipitates from HEK293 cells were shown for comparison. **B. Validation of circTau 12->7 protein using mass-spectrometry.** Immunoprecipitates were digested with trypsin and analyzed using mass-spectrometry. A representative spectrum shows peptide #7 that contains a methionine with an upstream sequence, indicating rolling circle translation. **C. Distribution of tau peptides on the mouse NP_001390913.1 reference sequence**. The reference sequence is 438 amino acids long. The part corresponding to the human circTau12->7 is indicated in gray. Peptide sequences are shown in Supplemental Figure 2. **D.** David analysis of circTau 12->7 interacting protein from mouse brain immunoprecipitates. Proteins are shown in Supplemental Figure 3. 167 proteins that co-immunoprecipitated with circTau 12->7 using anti-FLAG, but did not coimmunoprecipitated with circGFP or mRNA-GFP using anti-FLAG were clustered using DAVID with p<0.001.

### The circTau 12->7 RNA undergoes rolling circle translation and the resulting proteins interact with endogenous, mRNA encoded tau protein

To further characterize the protein made from the circTau 12->7 RNA in mouse brain, we performed mass spectrometry of the FLAG-immunoprecipitates from mouse brain generated from boiled lysates. Including phosphorylation variants, we observed 31 peptides (designated #2-28) covering the circTau 12-7 predicted peptide in the immunoprecipitates, which further confirmed the protein sequence predicted from the circTau 12->7 RNA (**Supplemental Figure 2**). Four of the peptides (#4, 10, 11 and 14) were specific for the human circTau 12->7 RNA, ruling out mouse mRNA encoded tau protein and demonstrating translation of the circTau RNA under physiological conditions. An uncomplete trypsin digest revealed the human-specific peptide #14 (LQTAPVPMPDLKNVK) that contained the only start codon in the circTau 12->7 RNA in the middle, not the start of the sequence, which demonstrated rolling circle translation. These finding were supported by the abundant peptide #15a that contained the only start codon (LQTAPVPMPDLK), but is also present in mouse. In the related peptide #15b, this start codon was oxidized at the methionine (LQTAPVPM*PDLK), indicating oxidative stress (22) (**Figure 3B**). As we previously reported (19), the circTau 12->7 encoded peptide interacts with endogenous, mRNA-encoded tau protein. Our mass spectrometry analysis confirmed these findings in mouse brain, as we detected five tau peptides outside the region encoded by circTau 12->7 in mouse brain immunoprecipitates (**Figure 3C**). In addition, we detected two mouse-specific peptides (#5, #26) in the region of the circTau12->7 encoded peptide.

Thus, the circTau 12->7 protein is translated in mouse through a rolling circle translation and the endoced protein co-aggregates with endogenous mouse mRNA encoded tau. This indicates that circTau12->7 endcoded protein causes aggregation of mRNA encoded tau protein.

### CircTau 12->7 protein interacts with proteins involved in vesicular transport

To identify other proteins that bind to circTau 12->7 encoded protein, we compared immunoprecipitates made from mouse brain injected with AAVs encoding circTau 12->7 circEGFP and mRNA-EGFP using mass-spectrometry. We considered only proteins as potential valid interactors that were present in the FLAG-immunoprecipitates from circTau 12->7 injected animals, both male and female, but absent in parallel immunoprecipitates from tissue infected with AAVs expression circEGFP or mRNA-EGFP. The presumed interacting proteins are listed in **Supplemental Figure 3**.

We considered as interacting proteins those that were represented in the mass spectrometry analyses by more than 10 independent peptides, were absent in IPs made from mice injected with AAVs expressing circGFP or mRNA-encoded GFP.

DAVID analysis (23) revealed an enrichment of proteins (p<0.001) involved in calmodulin binding (CAMKV, CAMK2B, CAMK2C), actin binding (SPTAN1, DBN1), myosins (Myo5a, Myo18a, Myo6), motor proteins (DNM1, DNM2) and numerous kinases. These data indicated a possible function of circTau 12->7 encoded peptide in affecting the actin cytoskeleton and/or intracellular transport. Kinases and calmodulin binding proteins were enriched in the anti-FLAG immunoprecipitants after circTau 12->7 AAV injections, suggesting a possible role of this complex in protein modifications and calcium sensing. It remains to be determined which of these interactions are due to a pathophysiological aggregation of the circTau 12->7 encoded protein (**Figure 3D).**

### CircTau 12->7 RNA is translated only in neurons

We next analyzed the translation of circTau 12->7 RNA on a cellular level, using immunohistochemistry. Following AAV circTau 12->7 infection, induced protein staining appears only in neurons, which we also observed for circEGFP. This was unexpected, since the circTau 12->7 construct is driven by the CAG promoter that is active in most known brain cells, including glia cells. We confirmed the neuronal staining using double-label immunofluorescence staining with NeuN as a neuronal marker. The double staining revealed coexpression between NeuN and circTau 12->7 and circEGFP, further confirming the neuron-specificity of the staining (**Figure 4A**). We next performed the similar experiment following viral infection with circEGFP. Again, we found a strong colocalization between the circularized protein and NeuN (**Figure 4B)**, indicating translation of circEGFP that is highly enriched in neurons .

**Figure 4.**
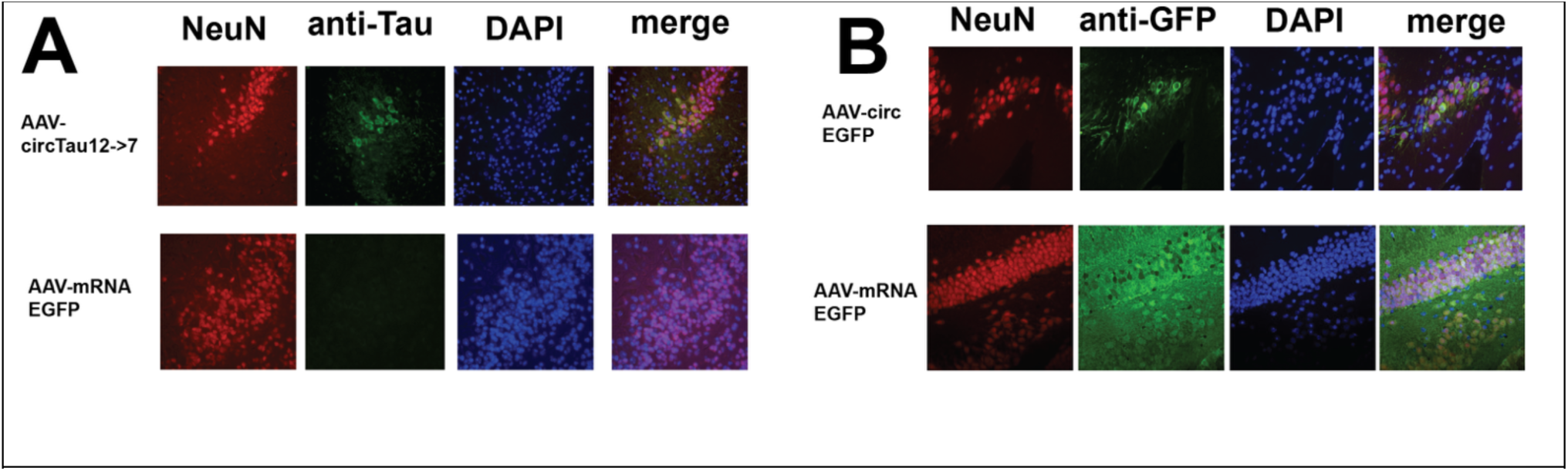
CircRNA, including circTau 12->7 RNA is translated in neurons. Expression of circular RNA and mRNA encoded proteins. **A:** Mice were injected with AAVs expressing circTau 12->7 or mRNA EGFP and **B:** were injected with AAVs expressing circEGFP and mRNA-EGPP. Hippocampal brain sections were stained with NeuN, anti-Tau, anti GFP and DAPI as indicated.

### CircTau 12->7 protein expressed in mouse brain from AAV infections caused aggregation of endogenous mRNA encoded tau protein into neurofibrillary tangle (NFT)-like structures

Since circTau 12->7 encoded protein interacts with mRNA-encoded endogenous tau protein (**Figure 3C**), we analyzed its intracellular distribution in circTau 12->7 AAV infected brain regions. Using C57BL/6 wildtype mice, 10^7^ GC (genomic copies) of the AAV containing pAAV circTau 12->7 were injected into the hippocampus. After 21 days, animals were fixed with paraformaldehyde via cardiac perfusion and the brains were embedded in paraffin. Microtome-cut brain sections were stained with anti-FLAG antibody to show the cell staining pattern. Using these methods, circTau 12->7 protein was localized in cornu ammonis neurons of the hippocampus, concentrated in the cell bodies and extending to the proximal neurites, with limited nuclear staining. We next performed immunohistochemical staining using a polyclonal anti-tau (nonphosphorylated) chicken antiserum that labels tau pathology in human AD neuropathologic changes (24) . We observed tau pathology in neurons, which was further confirmed when we used an antiserum against PHF-tau (phosphorylated Tau [pTau]) (**Figure 5A**) on adjacent sections.

**Figure 5.**
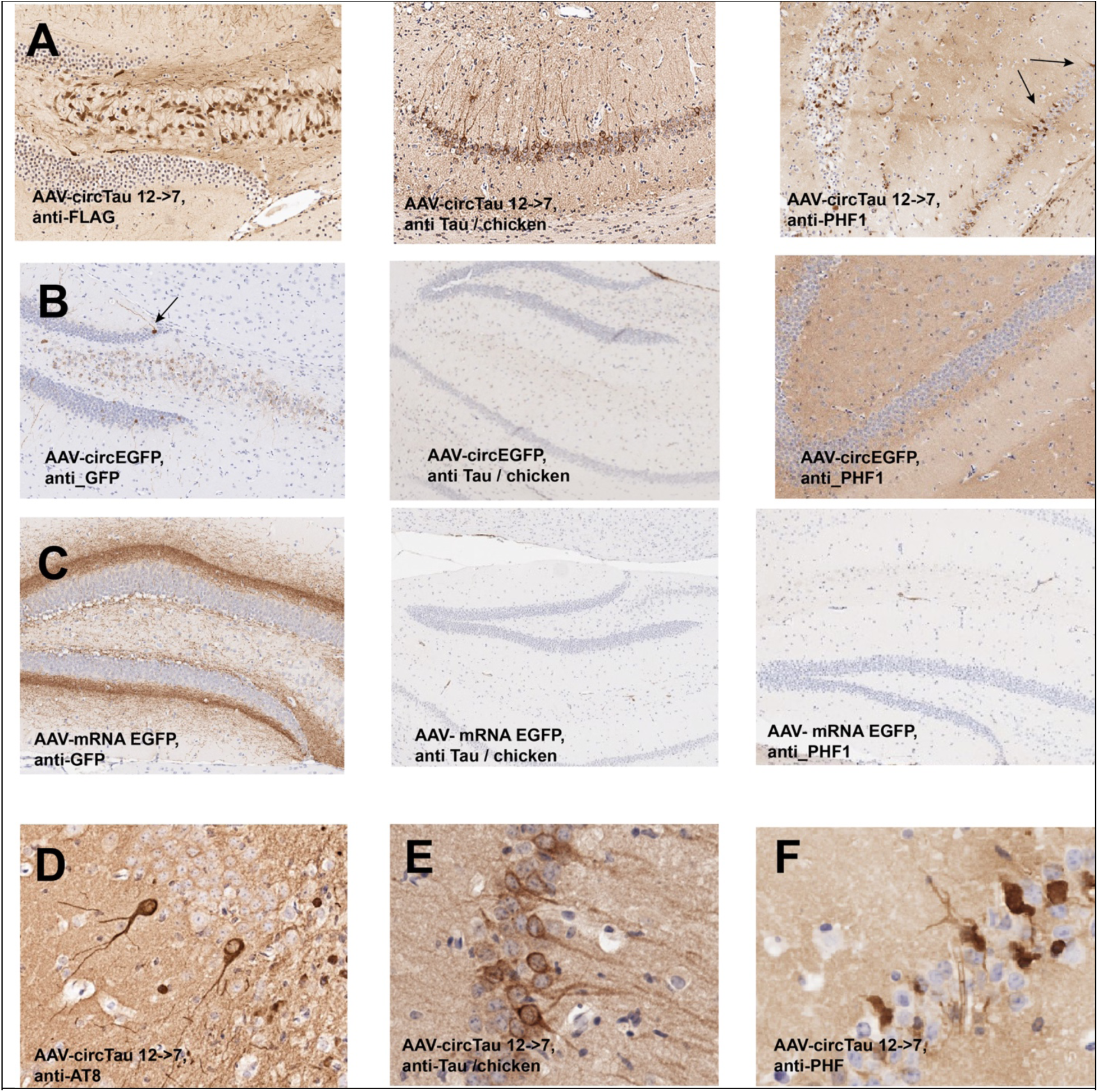
CircTau 12->7 RNA is translated in neurons and leads to NFT formation. Mice injected with Adeno associated viruses expressing the circRNAs indicated. Mice were fixed with paraformaldehyde and sections stained with the antibodies indicated, which were detected by ABC-DAB staining. Parallel sections on the right were stained with chicken anti-tau and an antibody against paired helical filaments. **A.** Injection with circTau 12-7 expressing AAV shows expression in hippocampal cornu ammonis neurons. **B.** Mice injected with AAV expressing circular EGFP were stained with a polyclonal anti GFP antibody and parallel sections **C.** Mice were injected with an AAV expressing mRNA-encoded EGFP. **D.** Sections of mice injected with AAV expressing circTau 12-7 were stained with anti phospho-tau (AT8). **E.** Enlargement of sections stained with anti-tau/chicken, and **F.** anti-PHF.

We employed two EGFP constructs as controls. To control for effects from possible circRNA translation, we generated an AAV expressing a circRNA encoding EGFP. The EGFP sequence was split after the tyrosine in the chromophore TLTYGVQ region (25), which ensures that green fluorescence is only generated after circularization (**Supplemental Data 4**). Injection of this virus into mouse hippocampus resulted in green fluorescence of a few hippocampal neurons. We did not observe tau pathology detected by the chicken tau antiserum serum or pTau antibody (**Figure 5B**). The second control was a commercial AAV with C9 serotype expressing mRNA encoded EGFP (Vectorbuilder, Chicago). Injection of this construct resulted in strong EGFP expression throughout the mouse brain detected by Western blot and intense staining via immunohistochemistry. Tau neuropathology could not be detected in these brains (**Figure 5C**).

Sections of mice injected with AAV expressing circTau 12-7 were stained with chicken anti-Tau antiserum and monoclonal pTau (AT8). As opposed to the findings in the control brains, the brains with AAV circTau transduction showed immunoreactive structures that closely resembled NFTs **(Figure 5D-F)**.

In summary, circTau 12->7 and possibly other circRNAs are translated predominantly in neurons. CircTau12->7 encoded protein promotes the formation of NFTs by causing the aggregation of mRNA encoded, endogenous tau protein.

## Discussion

In the present study, we evaluated circTau proteins expressed in mice brains. Following viral transduction, circTau proteins were detected preferentially in mouse neurons. The translation of circTau in mice brains resulted in peptides that were reminiscent of those from human cultured cells produced through rolling circle translation. Mass spectrometry assessments following mouse brain co-immunoprecipitation showed tau protein fragments from outside the AAV vector that derive from endogenous mouse tau protein were bound to circTau protein along with proteins involved in traffic and actin binding. These findings indicated that circTau can be expressed in mouse neurons, where the circRNA-derived can seed tau pathology that includes mRNA-translated full-length tau. It is possible that circTau 12->7 encoded protein fine-tunes vesicular traffic in neurons.

### CircTau 12->7 can be expressed in mouse brain

CircRNAs depend on secondary structures in the pre-mRNA to bring the backsplice sites together. In primates, these secondary structures are often imposed by Alu elements that comprise about 12% of the human genome, but that are missing in rodents. The MAPT gene contains at least 89 Alu elements that likely contribute to the human-specific expression of circTau RNAs (26). To recapitulate the conditions in human cells, we flanked the circTau 12->7 expression cassette with heterologous Alu elements that promoted the formation of circular RNA, both in human cells and in mouse brain. We observed the correct backsplicing in mice brains, suggesting that mouse cells recognize pre-mRNA structures mediated by Alu-elements.

### CircTau 12->7 is translated in mouse neurons

The circTau12->7 RNA was translated into protein both in human and in mouse cells. Confirming previous results where we used the authentic tau-Alu elements, ADAR activity promoted translation in HEK293 cells. Consisting with the increase in translation, we saw a shift in polysomes from the 80S initiation complex towards high molecular weight, actively translating polysomes. This observation is consistent with a model where ribosomes are initially stalled at the circular RNA. RNA editing releases this repression, possibly by opening secondary structures in the circRNA. We observe multiple protein bands that in size roughly correlate with a peptide generated from one round of rolling circle translation. Since there is no stop codon, the release of the peptides could be caused by ribosome collisions at the initiation site.

Three lines of evidence support that the proteins are made from a circular RNA, not from RNAs made by trans-splicing that have been observed in drosophila (27). First, the RNA made from the expression constructs is RNase R resistant, the biochemical criterion for a circular RNA. Secondly, we observe four different sizes of proteins generated from our expression constructs, which would indicate trans-splicing of four pre-mRNA, which has never been observed and is highly unlikely. Thirdly, we identified a peptide that contains amino acids downstream of the only initiation methionine, which supports a rolling circle, but did not found a peptide starting with a methionine.

### circTau RNAs as contributors for NFT formation

Using immunohistochemistry, we observed phosphor-Tau immunoreactivity and the typical morphology of neurofibrillary tangles in mouse brain infected with AAVs expressing circTau 12->7, which has been reported earlier (28). Mass-spec analysis of the immunoprecipitates showed the presence of mRNA encoded mouse tau proteins in immunoprecipitates containing circTau 12->7 encoded protein, suggesting an interaction between circTau 12->7 and mRNA encoded proteins. This interaction is reminiscent of the binding of the K18 peptide, representing roughly one round of circTau 12->7 translation, with mRNA encoded tau protein, leading to tau aggregation. Since there is no endogenous circTau 12->7 RNA in mice, we had to overexpress the human circTau RNA that contained a FLAG tag for detection. Similarly, Alu elements are not present in mice, and the molecular changes caused by their introduction are not known.

A notable feature of the pathologic tau inclusion induced by circTau transduction was neuron cell type specificity, which is reminiscent of NFTs in human (AD) brains. In other words, the circTau 12->7 RNA translation occurred only in neurons, leading to wild-type NFTs in mouse brain upon circTau 12->7 expression. In conclusion, NFT-like structures could be induced by circTau 12->7 in vivo. This implies that removing circTau RNAs from human brains could be a therapeutic approach for tauopathies, including AD.

## Acknowledgments

This work was supported by R01AG095675 and NSF-BSF MCB-2221921.

## Supplemental Data 1

### Sequence of circTau 12-7 expression construct

HIPK3 introns are in small cases and grey, the first of the stop codons in possible non-AUG open reading frames are underlined. The circTau 12->7 backsplice cassette is in capital letters. The start codon in indicated, the 3x FLAG tag is highlighted in yellow. The circTau RNA has only one in frame start codon and no stop codon. The primers used for RT-PCR are indicted by an arrow and underlined, generating a 228 nt long amplicon.

Taaatgaattagcctcagcctctcaaagtgctaggattacagggatctatactacaaattgtgaatttgacccttaagagtttc ttcctgatatttaaaattgaaaaaaaaattgttgacattaatatttcttctttccttttttttcttttcctttttttttttttttttgcag GGGGCTGATGGTGACTACAAAGACCACGACGGTGATTACAAAGATCATGACATCG ATTACAAGGACGACGACGACAAAGATGACGATGACAAGAAAACGAAGATCGCCAC ACCGCGGGGAGCAGCCCCTCCAGGCCAGAAGGGCCAGGCCAACGCCACCAGGA TTCCAGCAAAAACCCCGCCCGCTCCAAAGACACCACCCAGCTCTGGTGAACCTCC AAAATCAGGGGATCGCAGCGGCTACAGCAGCCCCGGCTCCCCAGGCACTCCCGG CAGCCGCTCCCGCACCCCGTCCCTTCCAACCCCACCCACCCGGGAGCCCAAGAA GGTGGCAGTGGTCCGTACTCCACCCAAGTCGCCGTCTTCCGCCAAGAGCCGCCT GCAGACAGCCCCCGTGCCCATGCCAGACCTGAAGAATGTCAAGTCCAAGATCGGC TCCACTGAGAACCTGAAGCACCAGCCGGGAGGCGGGAAGGTGCAGATAATTAATA AGAAGCTGGATCTTAGCAACGTCCAGTCCAAGTGTGGCTCAAAGGATAATATCAAA CACGTCCCGGGAGGCGGCAGTGTGCAAATAGTCTACAAACCAGTTGACCTGAGCA AGGTGACCTCCAAGTGTGGCTCATTAGGCAACATCCATCATAAACCAGGAGGTGG CCAGGTGGAAGTAAAATCTGAGAAGCTTGACTTCAAGGACAGAGTCCAGTCGAAG ATTGGGTCCCTGGACAATATCACCCACGTCCCTGGCGGAGGAAATAAAAAG gtaggtaacccaactagttaagcacaatttgaaaattgtttcctttgagtgtcatgtcaatgcccaaaaagtttcagatatttgg atttgagatgctcaacctgtataaggattcagaaagttattctgattaatgattttaagattcagatatacaataatcccagcaa cttgggaggctgaggcaggagaatcttgaacccaggagatggaggttgcagtgagccgagatcatgccattgcactcca

Predicted protein sequence of one round 27kDa MPDLKNVKSKIGSTENLKHQPGGGKVQIINKKLDLSNVQSKCGSKDNIKHVPGGGSVQIVYKPVDLSKVTSKCGSLGNIHHKPGGGQVEVKSEKLDFKDRVQSKIGSLDNITHVPGGGNKKGADGDYKDHDGDYKDHDIDYKDDDDKDDDDKKTKIATPRGAAPPGQKGQANATRIPAKTPPAPKTPPSSGEPPKSGDRSGYSSPGSPGTPGSRSRTPSLPTPPTREPKKVAVVRTPPKSPSSAKSRLQTAPVP

ATGCCAGACCTGAAGAATGTCAAGTCCAAGATCGGCTCCACTGAGAACCTGAAGCACCAGCCGGGAGGCGGGAAGGTGCAGATAATTAATAAGAAGCTGGATCTTAGCAACGTCCAGTCCAAGTGTGGCTCAAAGGATAATATCAAACACGTCCCGGGAGGCGGCAGTGTGCAAATAGTCTACAAACCAGTTGACCTGAGCAAGGTGACCTCCAAGTGTGGCTCATTAGGCAACATCCATCATAAACCAGGAGGTGGCCAGGTGGAAGTAAAATCTGAGAAGCTTGACTTCAAGGACAGAGTCCAGTCGAAGATTGGGTCCCTGGACAATATCACCC

ACGTCCCTGGCGGAGGAAATAAAAAGGGGGCTGATGGTGACTACAAAGACCACGACGGTGATTACAAAGATCATGACATCGATTACAAGGACGACGACGACAAAGATGACGATGACAAGAAAACGAAGATCGCCACACCGCGGGGAGCAGCCCCTCCAGGCCAGAAGGGCCAGGCCAACGCCACCAGGATTCCAGCAAAAACCCCGCCCGCTCCAAAGACACCACCCAGCTCTGGTGAACCTCCAAAATCAGGGGATCGCAGCGGCTACAGCAGCCCCGGCTCCCCAGGCACTCCCGGCAGCCGCTCCCGCACCCCGTCCCTTCCAACCCCACCCACCCGGGAGCCCAAGAAGGTGGCAGTGGTCCGTACTCCACCCAAGTCGCCGTCTTCCGCCAAGAGCCGCCTGCAGACAGCCCCCGTGCCC

## Supplemental Data 2

**A:** Mouse protein sequence for the MAPT reference gene (**NP_001390913.1),** length 438 aa. The sequence that corresponds to circTau 12-7 encoded protein in marked in gray. Peptides that are found both in mouse and human are marked in red. Peptides that are human-specific are marked in blue.

**B.** The sequence of the human circTau 12->7 encoded protein. The FLAG tag was omitted for clarification. Due to the high number of lysines, we did not observe peptides correponsing to the FLAG tag.

**A.** Mouse protein sequence (**NP_001390913.1)** MADPRQEFDTMEDHAGDYTLLQDQEGDMDHGLKAEEAGIGDTPNQEDQAAGHVTQARVASKDRTGNDEKKAKTSTPSCAKAPSHRPCLSPTRPTLGSSDPLIKPSSPAVCPEPATSPKHVSSVTPRNGSPGTKQMKLKGADGKTGAKIATPRGAASPAQKGTSNATRIPAKTTPSPKTPPGSGEPPKSGERSGYSSPGSPGTPGSRSRTPSLPTPPTREPKKVAVVRTPPKSPSASKSRLQTAPVPMPDLKNVRSKIGSTENLKHQPGGGKVQIINKKLDLSNVQSKCGSKDNIKHVPGGGSVQIVYKPVDLSKVTSKCGSLGNIHHKPGGGQVEVKSEKLDFKDRVQSKIGSLDN ITHVPGGGNKKIETHKLTFRENAKAKTDHGAEIVYK<u>SPVVSGDTSPR</u>HLSNVSSTGSIDM VDSPQLATLADEVSASLAKQGL

**B.** Human circTau 12->7 encoded sequence, 3x FLAG tag is underlined GADG<u>DYKDHDGDYKDHDIDYKDDDDKDDDDK</u>KTKIATPRGAAPPGQKGQANATRIPA KTPPAPKTPPSSGEPPKSGDRSGYSSPGSPGTPGSR<u>SRTPSLPTPPTREPK</u>KVAVVRTPPKS PSSAKSRLQTAPVPMPDLKNVKSKIGSTENLKHQPGGGKVQIINKKLDLSNVQSKCGSKDNIKHVPGGGSVQIVYKPVDLSKVTSKCGSLGNIHHKPGGGQVEVKSEKLDFKDRVQS K<u>IGSLDNITHVPGGGNKK</u>

## C. Table of observed peptide sequences

The peptides correspond to peptides present in the circTau 12-7 encoded protein and are marked in gray. # indicated a phosphorylation, location indicated the presence in Mouse protein sequence (**NP_001390913.1).** A # sign indicates phosphorylation, as * sign indicates oxidation.

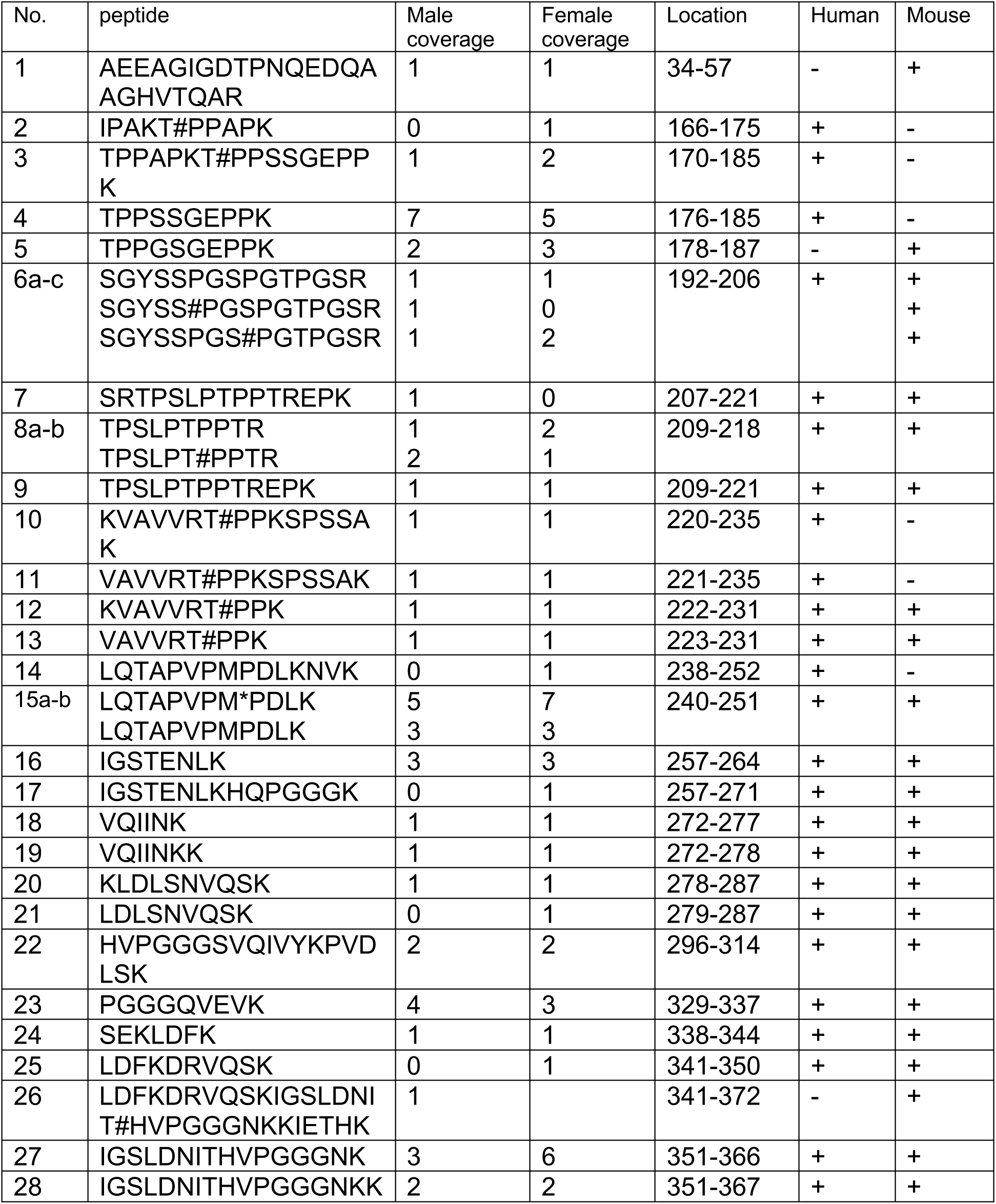
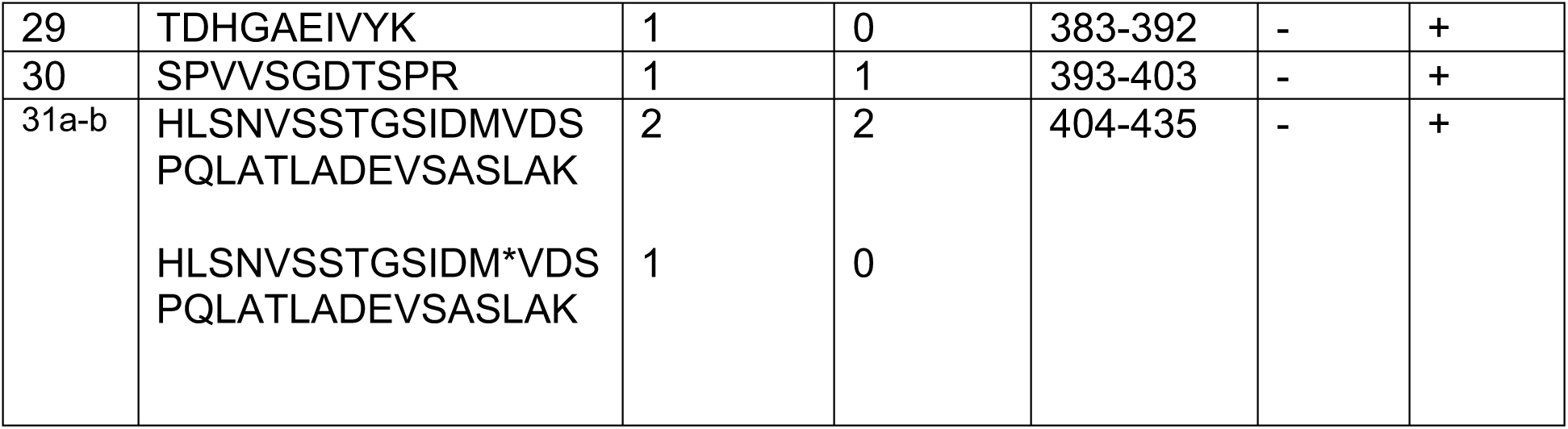

## Supplemental Data 3

### 12-7 Tau interacting proteins not found in EGFP immunoprecipitates

*Proteins detected after circTau 12-> injection and in neither circEGFP or mRNA EGFP injection*

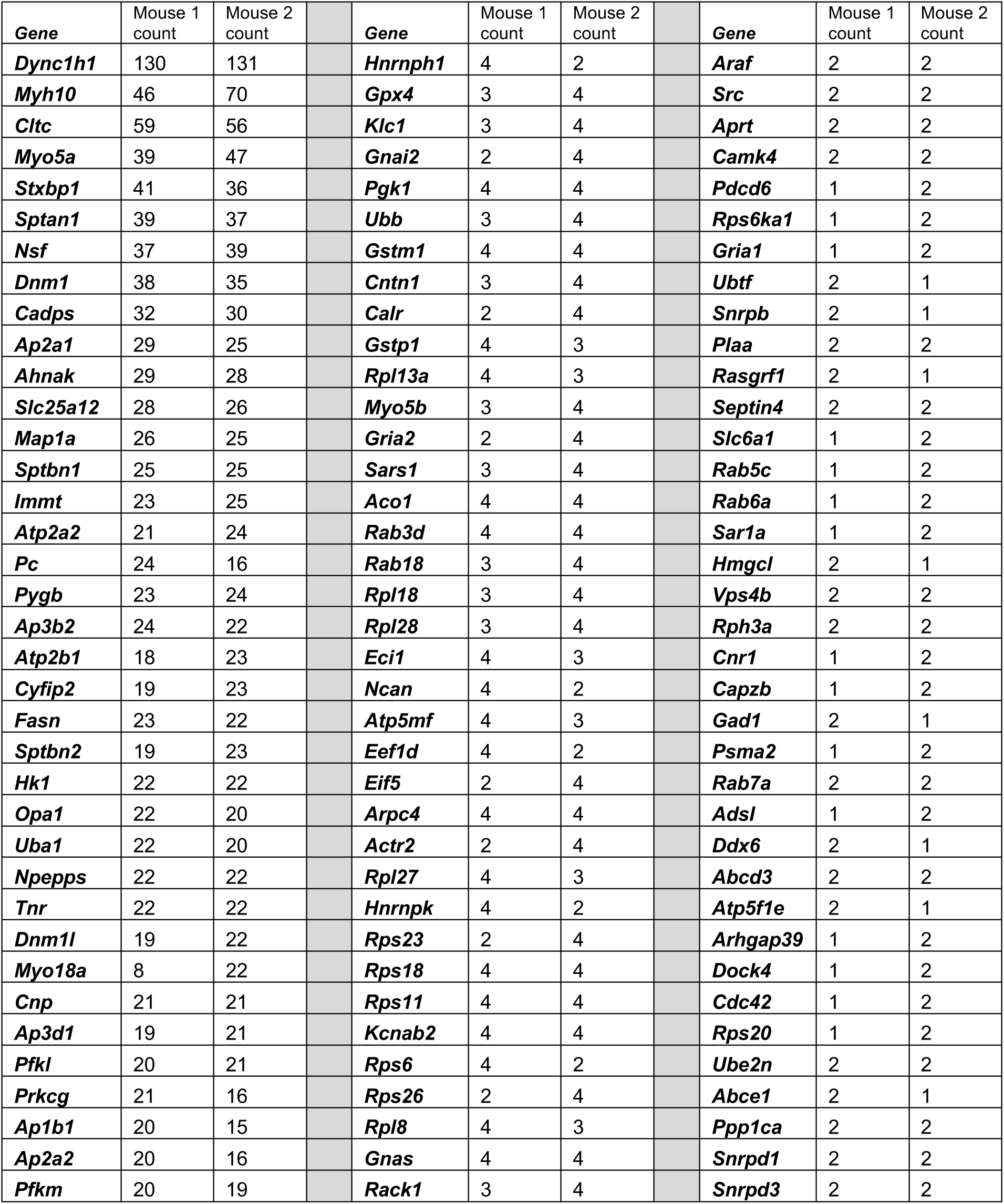
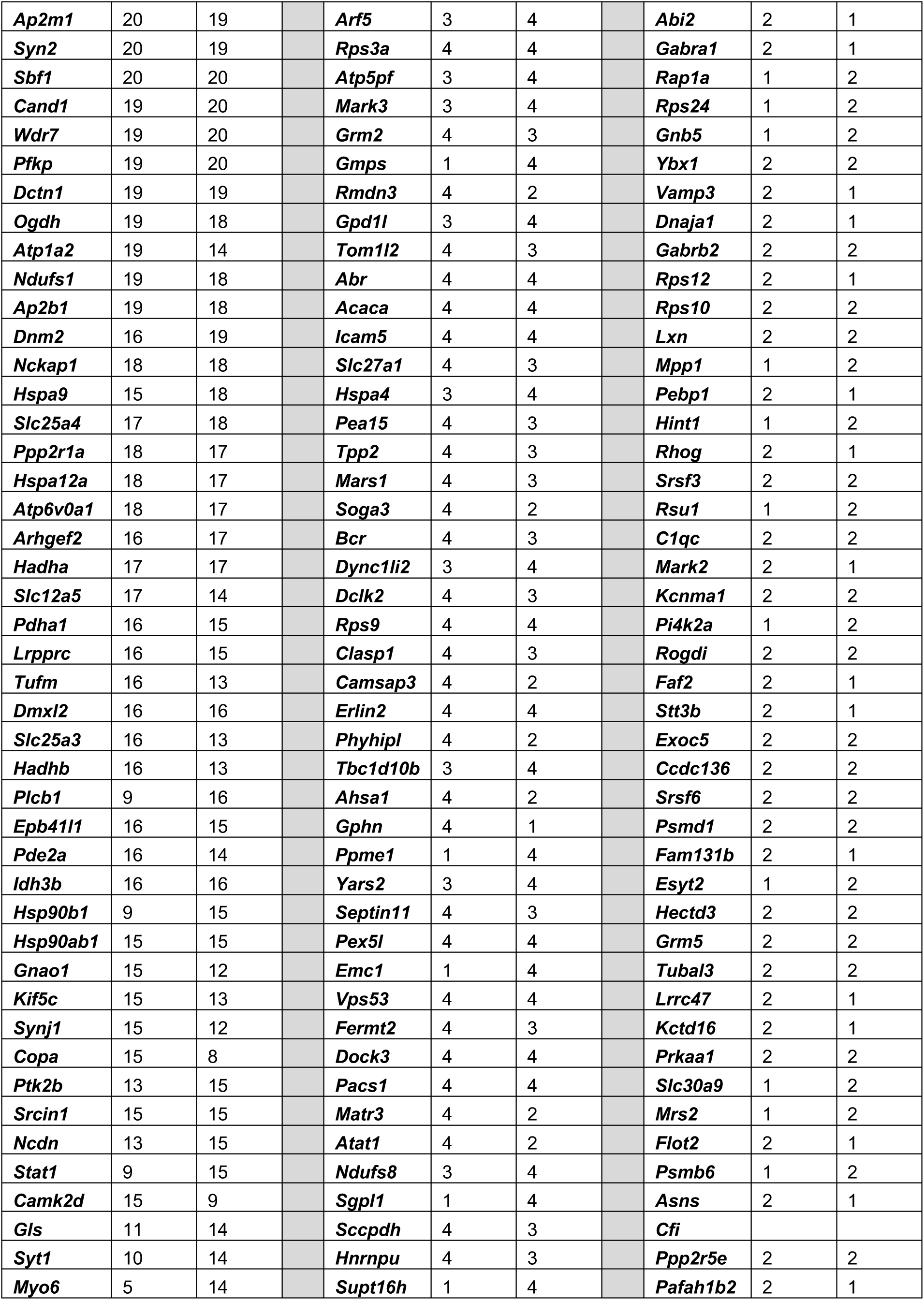
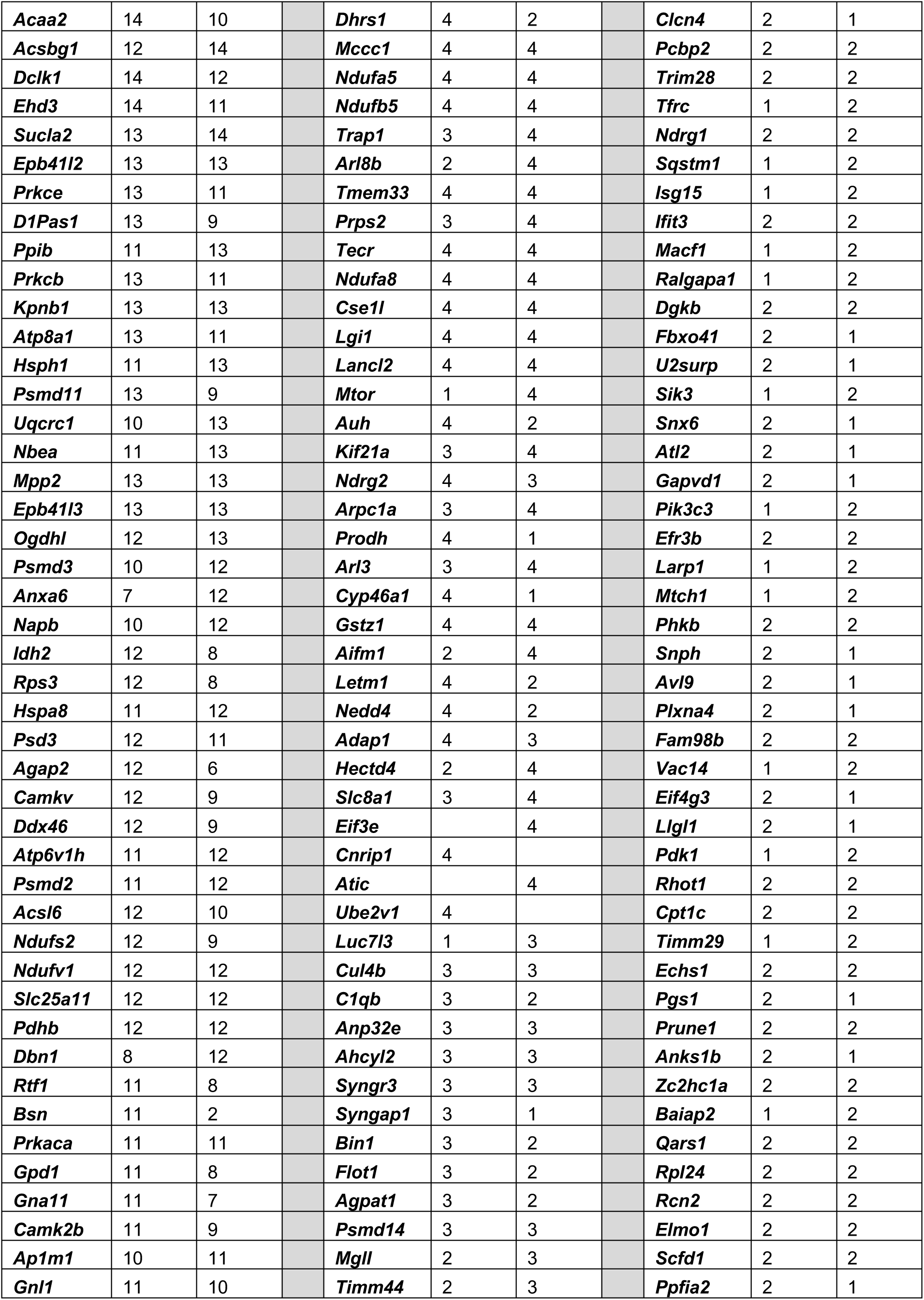
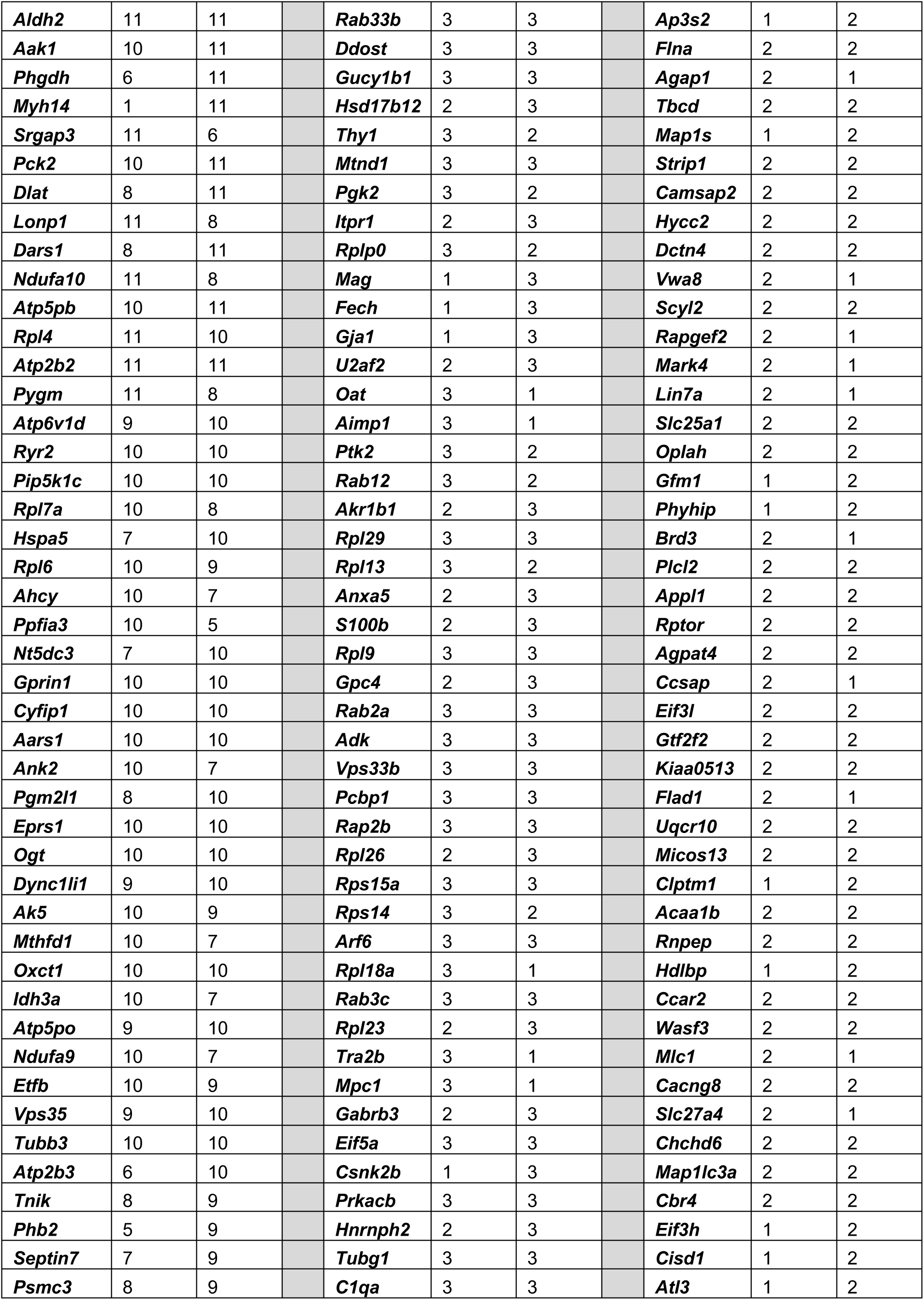
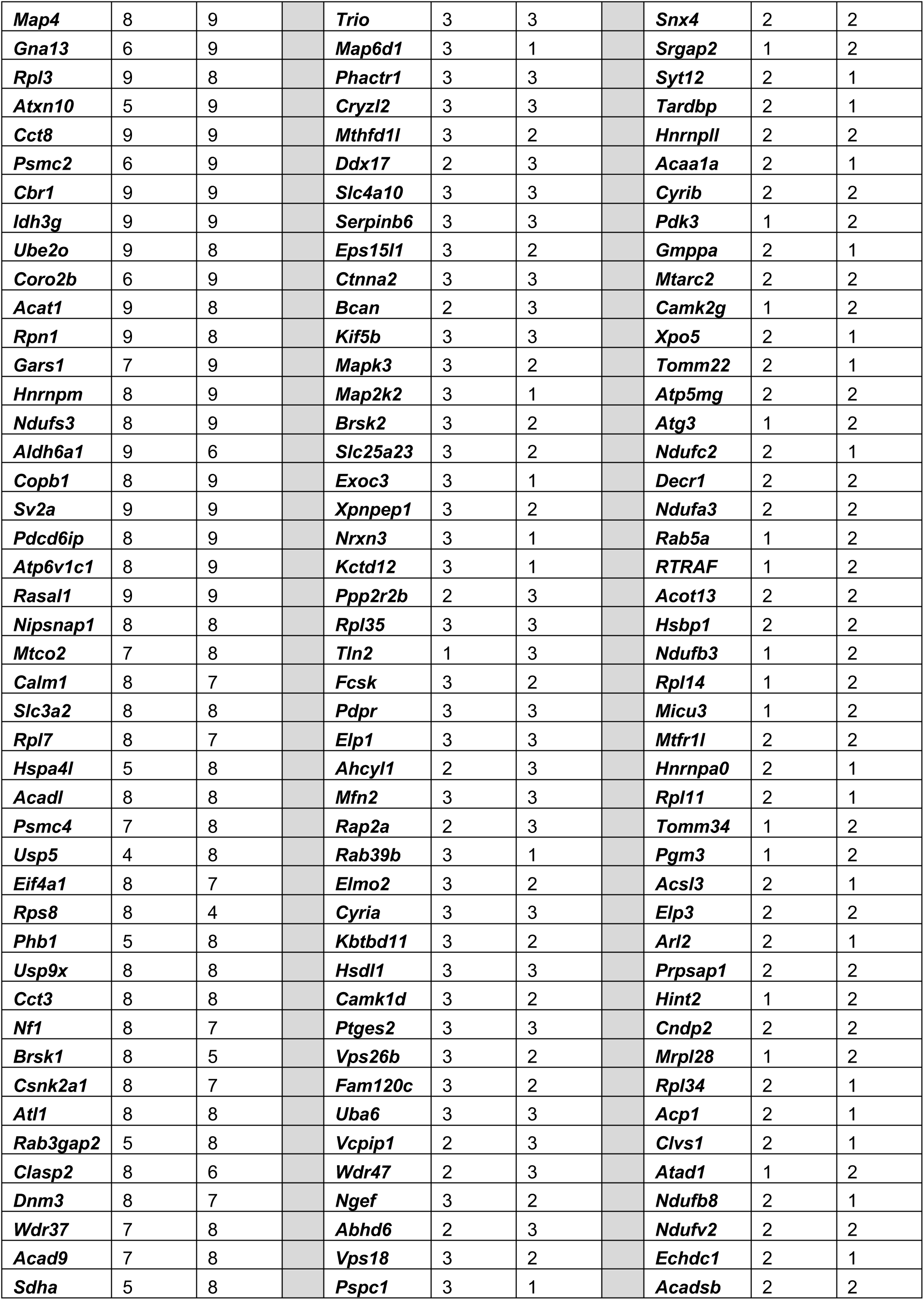
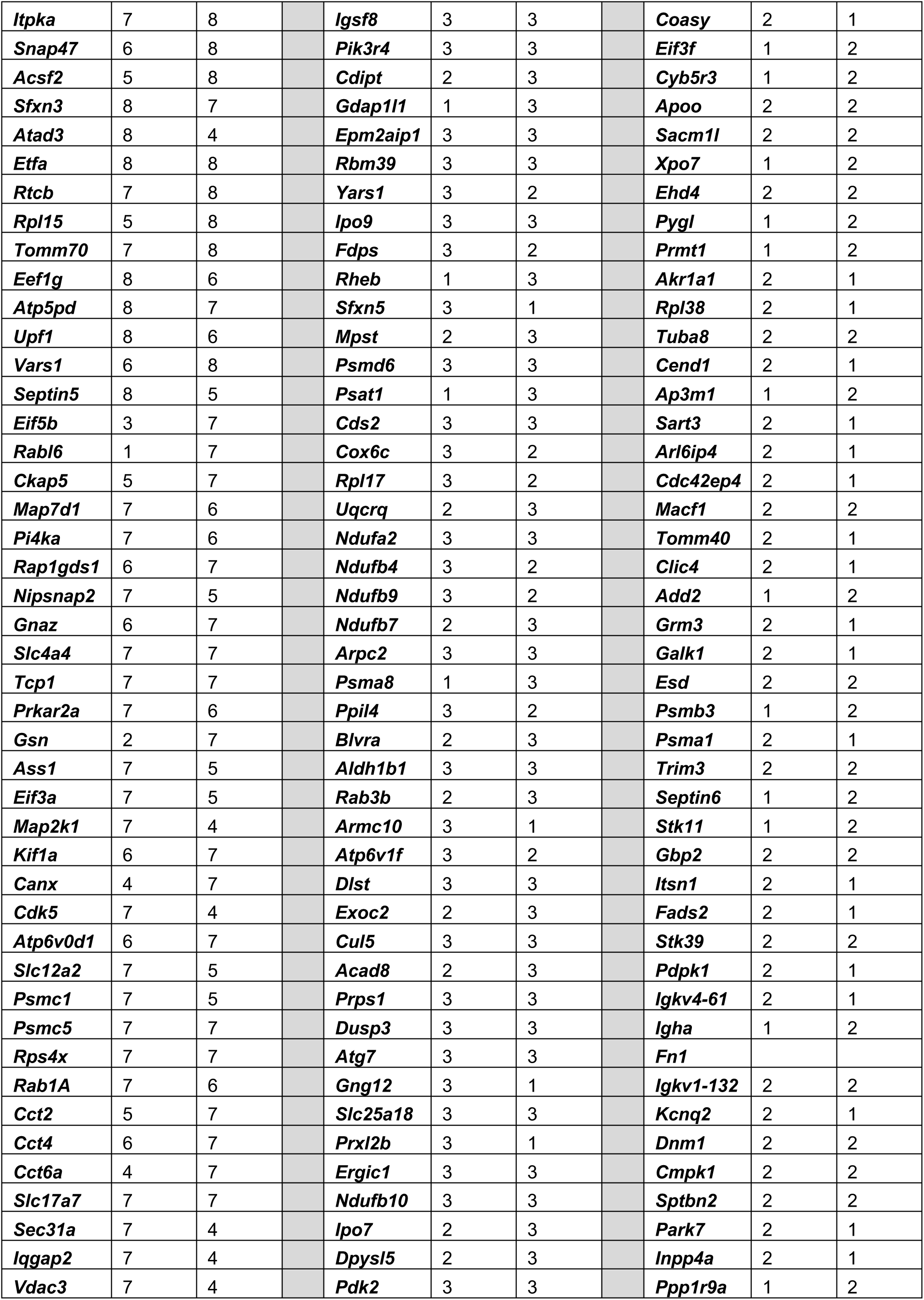
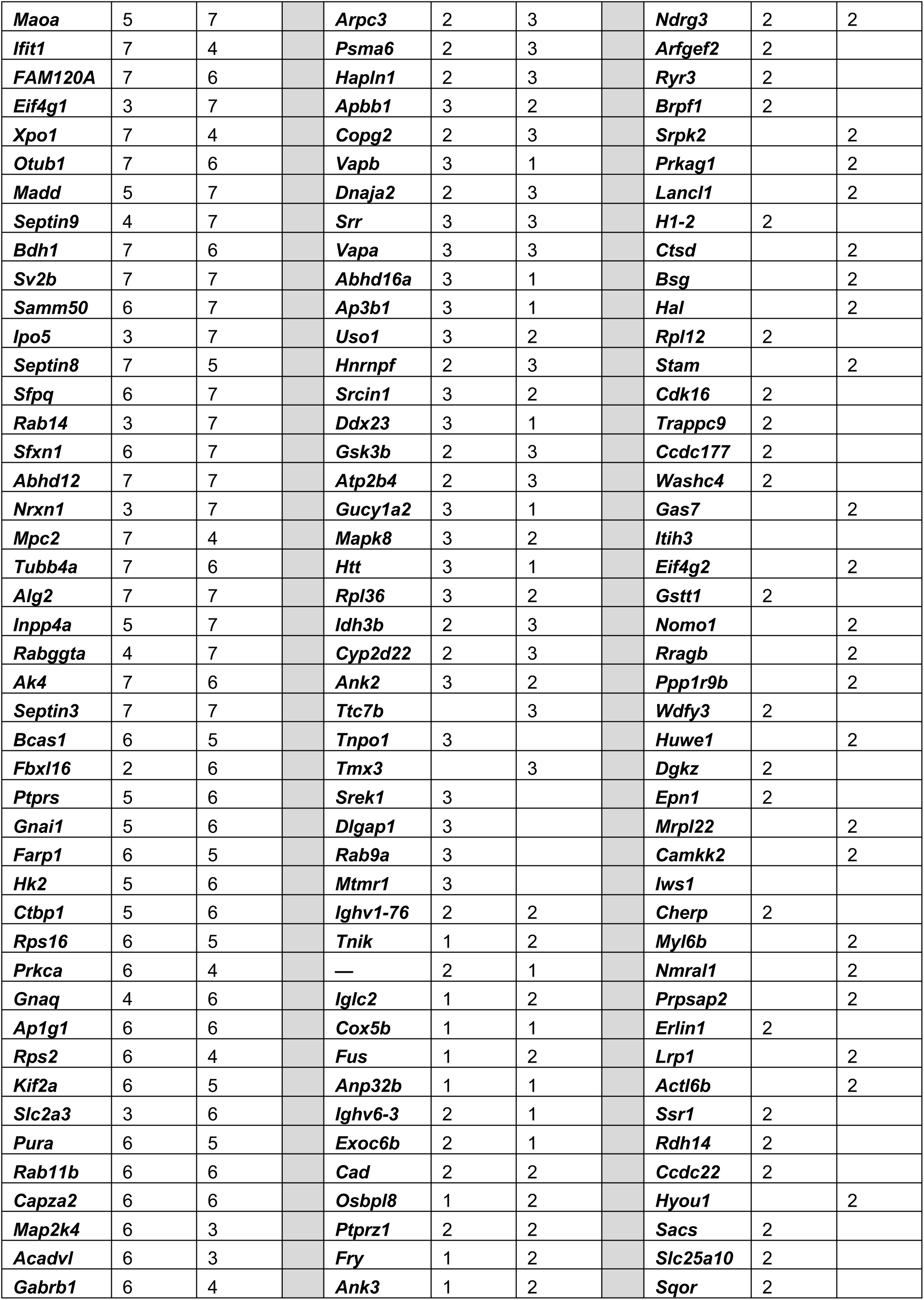
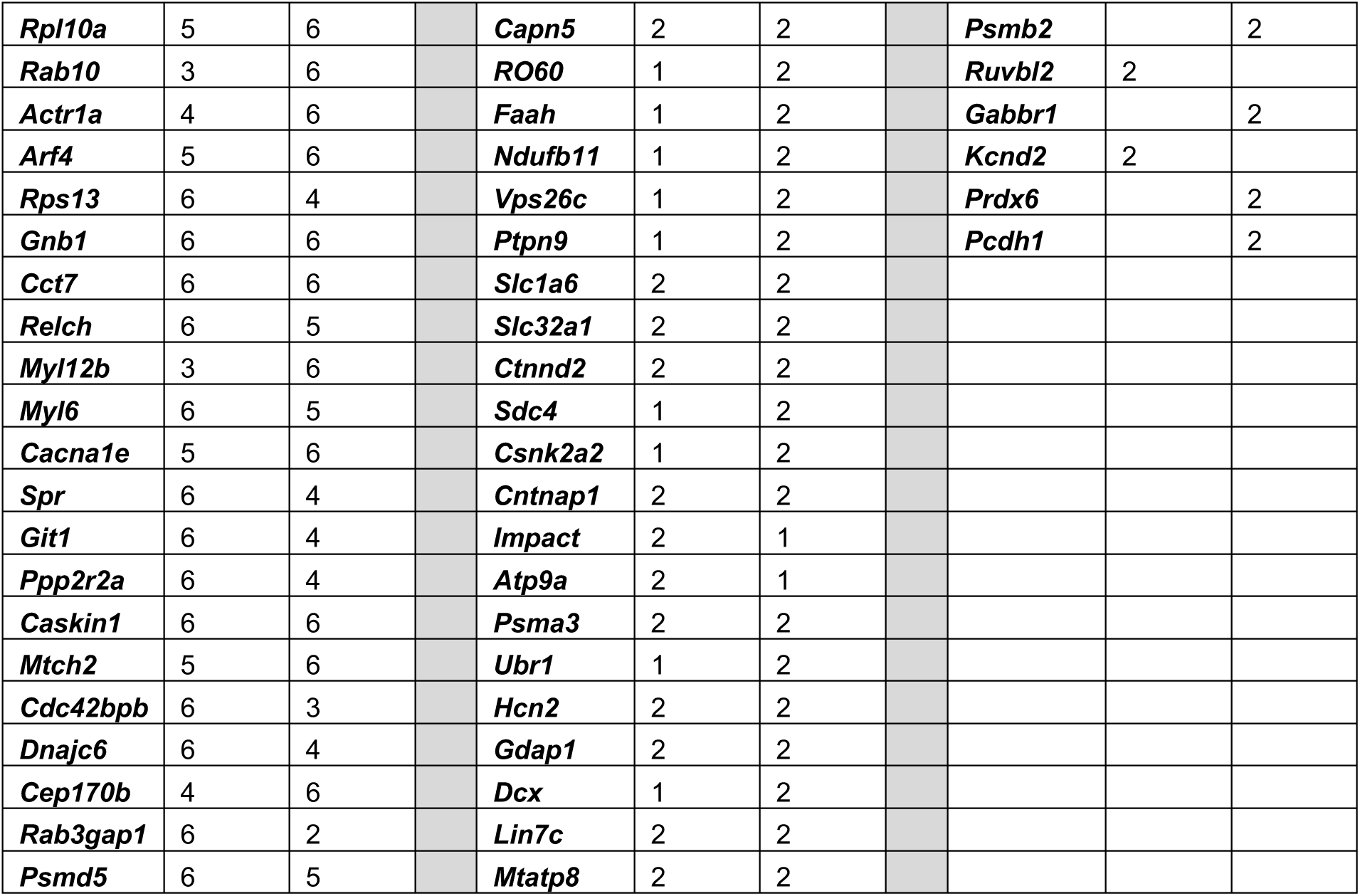

### Supplemental Data 4

Sequence of the circEGFP construct

The EGFP sequence was splice after the tyrosine in the chromophore TLTYGVQ, leaving the first G of the glycine codon at the 5’ backsplice site. This cassette was flanked by HIPK3 introns that were used to circularize the circTau 12->7 sequence. The underline sequence is a PP7 tag. The amino acids of the split Thr Leu Thr Tyr Gly Val Gln chromophore are indicated.

Taaaagaattagcctcagcctctcaaagtgctaggattacagggatctatactacaaattgtgaatttgacccttaagagtttcttcctgatatttaaaattgaaaaaaaaattgttgacattaatatttcttctttccttttttttcttttcctttttttttttttttttgcag GCGTGCAGTGCTTCAGCCGCTACCCCGACCACATGAAGCAGCACGACTTCTTCAAGTCCGCCATGlyValGln CCCGAAGGCTACGTCCAGGAGCGCACCATCTTCTTCAAGGACGACGGCAACTACAAGACCCGCGCCGAGGTGAAGTTCGAGGGCGACACCCTGGTGAACCGCATCGAGCTGAAGGGCATCGACTTCAAGGAGGACGGCAACATCCTGGGGCACAAGCTGGAGTACAACTACAACAGCCACAACGTCTATATCATGGCCGACAAGCAGAAGAACGGCATCAAGGTGAACTTCAAGATCCGCCACAACATCGAGGACGGCAGCGTGCAGCTCGCCGACCACTACCAGCAGAACACCCCCATCGGCGACGGCCCCGTGCTGCTGCCCGACAACCACTACCTGAGCACCCAGTCCGCCCTGAGCAAAGACCCCAACGAGAAGCGCGATCACATGGTCCTGCTGGAGTTCGTGACCGCCGCCGGGATCACTCTCGGCATGGACGAGCTGTACAAGTCCGGCCGGACTCAGATCTCGAGCTCAAGCTTCGTGCAGAACCAGCAGAGCATATGGGCTCGCTGGCTGCACGTCGACTCTAGAAACCAGCAGAGCATATGGGCTCGCTGGTAGccaccATGGTGAGCAAGGGCGAGGAGCTGTTCACCGGGGTGGTGCCCATCCTGGTCGAGCTGGACGGCGACGTAAACGGCCACAAGTTCAGCGTGTCCGGCGAGGGCGAGGGCGATGCCACCTACGGCAAGCTGACCCTGAAGTTCATCTGCACCACCGGCAAGCTGCCCGTGCCCTGGCCCACCCTCGTGACC ACC CTG ACC TAC G

Thr Leu Thr Tyr G gtaggtaacccaactagttaagcacaatttgaaaattgtttcctttgagtgtcatgtcaatgccc aaaaagtttcagatatttggatttgagatgctcaacctgtataaggattcagaaagttattctga ttaatgattttaagattcagatatacaataatcccagcaacttgggaggctgaggcaggagaatc ttgaacccaggagatggaggttgcagtgagccgagatcatgccattgcactcca

